# A method of back-calculating the log odds ratio and standard error of the log odds ratio from the reported group-level risk of disease

**DOI:** 10.1101/760942

**Authors:** Dapeng Hu, Chong Wang, Annette M O’Connor

## Abstract

In clinical trials and observational studies, the effect of an intervention or exposure can be reported as an absolute or relative comparative measure such as risk difference, odds ratio or risk ratio, or at the group level with the estimated risk of disease in each group. For meta-analysis of results with covariate adjustment, the log of the odds ratio (log odds ratio), with its standard error, is a commonly used measure of effect. However, extracting the adjusted log odds ratio from the reported estimates of disease risk in each group is not straightforward. Here, we propose a method to transform the adjusted probability of the event in each group to the log of the odds ratio and obtain the appropriate (approximate) standard error, which can then use used in a meta-analysis. We also use example data to compare our method with two other methods and show that our method is superior in calculating the standard error of the log odds ratio.

## Background

Many randomized controlled trials (RCT) are conducted in livestock production facilities. In these studies, although the animals are individually randomized to treatment group, the treatments groups are housed together in units. These housing units can be referred to variably as pens, houses, barns, rooms, pastures, villages, etc. and differ in name based on the species and country. Regardless of the name, housing is associated with a commonality of ration and environment that very often impacts important outcomes such as mortality, morbidity and growth. In these livestock populations, the housing unit is therefore considered a source of dependency in the outcome. To account for this dependency, it is often recommended that a random effect for the housing unit be incorporated into the model estimating the intervention effect size. Similarly, in observational studies, dependency in the outcome of animals housed together is commonly adjusted for by inclusion of a random effect for housing (pen, barn, house, etc.) [1] [2] [3] [4]. This dependency in the outcome is sometimes also called clustering and readers interested in a more detail explanation of dependency of outcomes in livestock populations are directed to other publications [2].

The need to adjust for housing effects has implications for the reporting of study effects. For example, the REFLECT statement, which is a guideline for reporting clinical trials in livestock production, suggests that authors report *“For each primary and secondary outcome, a summary of results for each group, accounting for each relevant level of the organizational structure, and the estimated effect size and its precision (e.g. 95% confidence interval)”*. The rationale for requesting group-level risk information, in additional to relative estimates, is that risk is easier to interpret for end users, compared to adjusted relative estimates, which are more useful to meta-analysis. Group-level estimates of disease risk can be obtained using standard statistical analysis software. For example, in the `GLIMMIX` procedure from SAS (version 14.1), the group-level estimates of disease risk are obtained by applying the inverse link function to the least-squares means estimates reported in the outcome [5]. For studies that use random effects models, the effect sizes should all be adjusted estimates.

Interestingly, in a recent review of veterinary clinical trials we noted that several authors of clinical trials in livestock production chose to report only the adjusted group-level estimates of disease risk and corresponding 95% confidence intervals [6] [7], not a relative effect size (odds ratio or risk ratio). To illustrate, Schunicht et al in a clinical trial of antibiotics to control undifferentiated fever in feedlot cattle reported using the following approach to analysis *The animal health variables were compared between the experimental groups by using linear logistic regression modeling techniques controlling for within-pen clustering of disease by using the method previously described and reviewed (25,26)* [6]. The authors then reported the initial undifferentiated fever risk as 23.17% for the oxytetrcycline group and 18.32% for the tilmicosin group with a common standard error of 1.59% . The authors also reported the p-value of 0.046 for the treatment effect, however the odds ratio was not explicitly reported [6].

For meta-analysis of adjusted effect sizes the log odds ratio is often used. When authors only report the adjusted disease risk per group it is necessary to convert the group-level risk back to a log of the odds ratio (also called the log odds ratio). However, we were unable to find guidance for converting the risk estimates to the log odds ratios in standard meta-analysis texts [8] [9] [10]. In personal communications with researchers who work on similar topics and have encountered the same issue, it appears that at least two approaches have been used. We call these naive approaches because they do not appear to be based on any proposed published methods. Here, we propose an approach to obtaining estimates of effect size (log odds ratio) and its standard error (SE) using only the point estimate of the risk of disease in each treatment group and the significance test results (p-value) of the treatment effect size. We compare our proposed method to two naive approaches described to us by other review teams.

## Methods

In the following section, we first describe the random effects model on which we based our further analysis. We then propose a novel method to transform the adjusted group-level risk back to the log odds ratio averaged across random effects and present a method to approximate its standard error. The adjusted group-level risk is on a probability scale between 0 and 1; if a percentage is reported it can be converted to the probability scale first before apply our proposed method for transformation.

### The adjusted group-level risk obtained from a generalized linear mixed model(GLMM) for binomial outcome data

We consider a random effects model with two treatment groups, both shared with *J* blocks. Let *n*_*ij*_ be the number of experimental units that received treatment *i*(*i* = 1, 2) in block *j*(*j* = 1, …, *J*) and *r*_*ij*_ be the number of events. To compare the performance of these treatments and account for dependence, one can model the probability of treatment effect, *π*_*ij*_, by using the generalized linear mixed model with a logit link function. The model is:

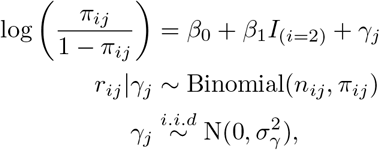

where *γ*_*j*_ denotes the random block effects of block *j* and 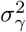 is the variance of the random block effects. *I* is the indicator function.

Different estimation methods have been proposed to maximize the likelihood function of the GLMM. In the `lme4`, `Zelig` and `glmmML` packages from R (version 3.5.2), the estimation methods that can be selected are the Laplace approximation developed by Daniels (1954) [11], the Barndorff-Nielsen and Cox approach (1979) [12], and the Gauss-Hermite quadrature approach. Monte Carlo likelihood approximation is used in the glmm package to find the Monte Carlo maximum likelihood estimates for the fixed effects and the variance components [13]. For testing the significance of any fixed treatment effects, the test statistic under the null hypothesis in these R packages approximates a normal distribution. In SAS 14.1, the default method of parameter estimation for models with random effects in `GLIMMIX` is the restricted pseudo-likelihood method as demonstrated by Wolfinger and O’Connell (1993) [14]. However, comparing these estimation methods is not the aim of this paper. Our goal is to document an approach to converting the adjusted group-level risk of the outcome to the adjusted log odds ratio averaged across the random effects estimate regardless of the estimation method used. For authors who use SAS for analyses, the adjusted group-level risk is often called the least squares means on the probability scale in the the default SAS outputs format. It is:

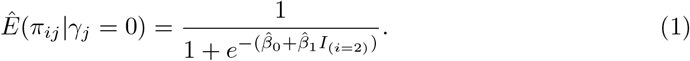

### Converting the adjusted disease risk back to a log odds ratio

For conducting meta-analysis, we need to transform estimated group-level risk, written here on the probability scale (0-1) *Ê*(*π*_*ij*_|*γ*_*j*_ = 0) back to a log odds ratio estimate 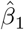 and its standard error. Suppose *Ê*(*π*_1*j*_|*γ*_*j*_ = 0) and *Ê*(*π*_2*j*_|*γ*_*j*_ = 0) were reported, then the point estimate of *β*_1_ is given by the standard formula:

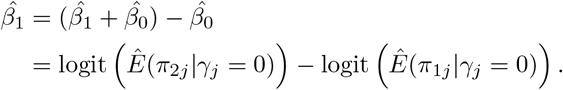

To obtain the standard error (SE) of 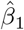, we use the p-value (*p*) for testing the significance of the fixed effect,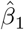 and assume the normality of the test statistic of *β*_1_. Then we have:

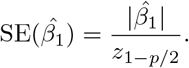

Note that for testing the significance of the fixed treatment effect in some software packages (`GLIMMIX` in SAS, `hglm` in R), the test statistic under the null hypothesis approximates a t-distribution. Although there are several methods available of determining the degree of freedom for the approximate t-distribution, containment is the default method in `GLIMMIX` if the model contains random effects. If degrees of freedom are reported in the original studies, then we can use t-distribution as the reference distribution.

For the case where there are *J* blocks within each treatment group *i*, the model is:

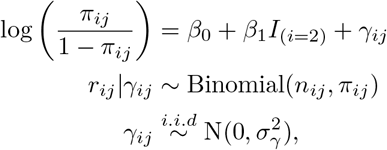

where *γ*_*ij*_ is the random block effects of block *j* within treatment *i*. The method proposed above also applies here.

### Two naive methods

As discussed, we were unable to find guidance as to how to transform the group-level adjusted risk, and therefore we evaluated the three possible approaches we have identified. The two naive ways are the relatively simple approaches suggested based on personal communications with other review authors. We present these for comparison’s sake; however, readers should be aware that we could find no citations recommending these approaches, hence we label them as naive. Suppose we have an RCT with two treatments *t*_1_ and *t*_2_. Let *n*_1_ and *n*_2_ denote the total number of enrolled animals in the two treatment groups. *p*_1_ and *p*_2_ denote the adjusted group-level disease risk calculated using equation 1, while *l*_1_ and *l*_2_ are the lower 95% confidence limits, and *u*_1_ and *u*_2_ are the upper 95% confidence limits. One suggestion was to transform the adjusted disease risk and the confidence intervals directly. Let 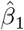 be the point estimate of the log odds ratio, which is obtained as follows:

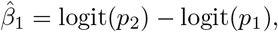

Then, the estimates of lower limits and upper limits of 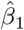 are obtained as follows

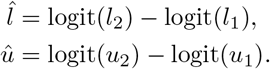

Then we can calculate the standard error of 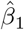 by dividing the range of the confidence interval by the 97.5% quantile of the standard normal distribution as follows:

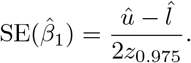

Another proposed method was to convert probabilities to raw frequency data based on the number of study units and then calculate log odds ratio and standard error using those frequency data. By this method,

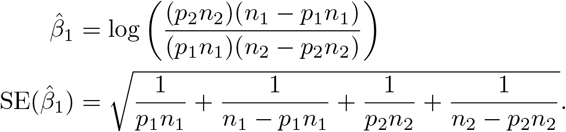

## Examples

Here, we provide examples of three possible conversion approaches using two data sets. The first data came from a 2-arm RCT conducted in swine. The objective of this study was to compare the efficacy of two interventions administered at the individual level in terms of the risk of a pig being treated or not. In this data, the random effects of different pens nested within different rooms create dependency that should be accounted for in this model. Table 1 shows the adjusted probability of being treated and the 95% confidence limits. The p-value of testing significant differences between the two groups is 0.0000299.

**Table 1.** Adjusted group-level risk of disease and 95% confidence intervals reported on the probability scale (0-1) for RCT data using results from a generalized linear model (`lme4` package). The model contains a fixed effect for treatment and a random effect for pens (n=24) nested within rooms (n=2)

| Treatment | Risk | Lower 96% Limit | Upper 95% Limits |
| --- | --- | --- | --- |
| Treatment A | 0.1939 | 0.1474 | 0.2507 |
| Treatment B | 0.2996 | 0.2381 | 0.3693 |

Table 2 shows the comparison of two naive methods with the method proposed and true estimation results. All three methods provide an accurate point estimate of log odds ratio, while our proposed method outperforms the naive methods in terms of estimation of the standard error. The difference in the estimate of the standard error for each method and the true standard error are also reported.

**Table 2.**
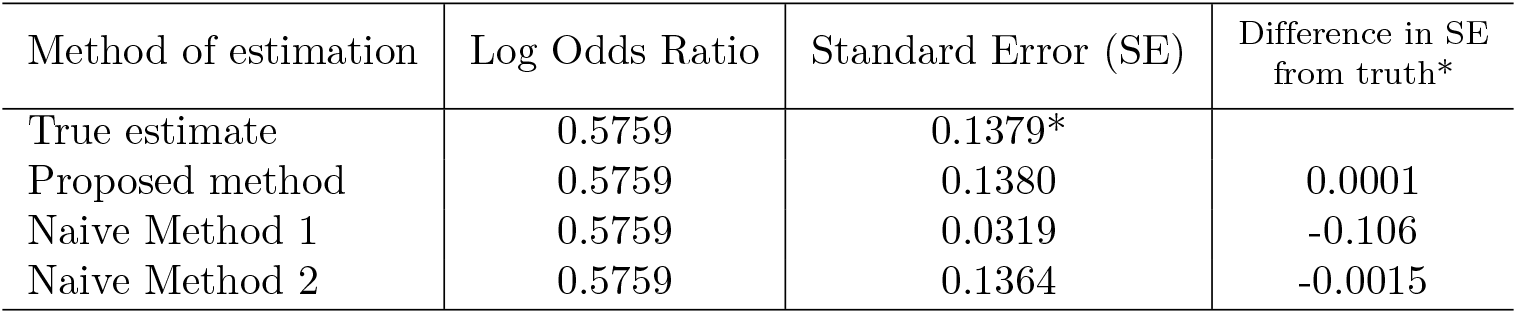
Estimates of Log odds ratio and its standard error corresponding to the RCT data in Table 1 using three methods of calculation

| Method of estimation | Log Odds Ratio | Standard Error (SE) | Difference in SE from truth* |
| --- | --- | --- | --- |
| True estimate | 0.5759 | 0.1379* |  |
| Proposed method | 0.5759 | 0.1380 | 0.0001 |
| Naive Method 1 | 0.5759 | 0.0319 | -0.106 |
| Naive Method 2 | 0.5759 | 0.1364 | -0.0015 |

Another example is an observational study and the data is a subset of the dataset “cbpp” from the R package lme4. The data description link is available at https://cran.r-project.org/web/packages/lme4/lme4.pdf here. The aim of these data are to study the within-herd spread of Contagious Bovine Pleuropneumonia (CBPP) at different time periods (period 1 and 2) in infected herds. In this example, cattle from multiple districts are studied, and therefore districts create dependency that should be considered as a random effect in this model, much the same way that housing units would create dependency in the swine RCT. Table 3 shows the adjusted disease risk of the period effect and the 95% confidence limits. The p-value of the period effect is 0.00127.

**Table 3.**
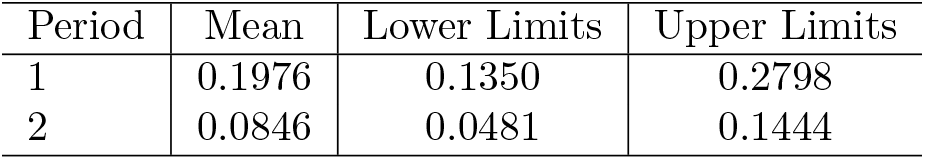
Adjusted group-level risk of disease and 95% confidence intervals reported on the probability scale (0-1) for observational data using results from a generalized linear model (`lme4` package). The model contains a fixed effect for period and a random effect for district (n=15)

As with the RCT example, this observational study (Table 4) shows that the point estimates of the log odds ratio are accurate for all approaches but the standard error of our proposed method has the smallest difference from the estimate given by the software results. The extent of the difference is smallest using our proposed approach and therefore we propose this method.

**Table 4.** Estimation of the Log odds ratio and its standard error comparison corresponding to the observational study data reported in Table 3

| Method of Estimation | Log Odds Ratio | Standard Error | Difference in SE from truth* |
| --- | --- | --- | --- |
| True estimate | -0.9807 | 0.3042* |  |
| Proposed method | -0.9807 | 0.3043 | 0.0001 |
| Naive Method 1 | -0.9807 | 0.2881 | -0.0161 |
| Naive Method 2 | -0.9807 | 0.2892 | -0.015 |

## Discussion

The aim of this paper is to help researchers extracting results from studies that only report the adjusted disease risk of each arm and the significance test results of the effect size and back-transform these to the log odds ratio averaged across a random effects scale. Our rationale for sharing this information is that this approach to reporting was more common than we expected in livestock reviews. For example, in one recent review, at least 10 of 75 relevant studies used this approach to reporting. Without a method to accurate estimate the log odds ratio and its standard error, the results of such studies might be excluded from systematic reviews and contribute to research wastage. We should note, that for parameter estimation in a generalized linear mixed model, different software has different estimation methods. These estimations methods although of interest, are not relevant here as our method seeks to convert the adjusted group-level disease risk reported on a probability scale (i.e., between 0 and 1) to the log odds ratio no matter what estimation method is used. One potential limitations of our approach is requirement that the p-value of the treatment is reported. If a study contains more than two treatments, this method is also valid if corresponding p-values are reported.

## References

1. Sargeant J, O’Connor AM, Dohoo I, Erb H, Cevallos M, Egger M, et al. Methods and processes of developing the Strengthening the Reporting of Observational Studies in Epidemiology–Veterinary (STROBE-Vet) statement. Journal of veterinary internal medicine. 2016;30(6):1887–1895.

2. O’Connor AM, Sargeant J, Dohoo I, Erb H, Cevallos M, Egger M, et al. Explanation and elaboration document for the STROBE-vet statement: Strengthening the reporting of observational studies in epidemiology–veterinary extension. Zoonoses and public health. 2016;63(8):662–698.

3. O’Connor AM, Sargeant JM, Gardner IA, Dickson JS, Torrence ME, consensus meeting participants: CE Dewey REJGMGGKSLPMARWSDSKSJSMWRW IR Dohoo. The REFLECT statement: methods and processes of creating reporting guidelines for randomized controlled trials for livestock and food safety. Journal of veterinary internal medicine. 2010;24(1):57–64.

4. Sargeant JM, O’Connor AM, Gardner IA, Dickson JS, Torrence ME, Dohoo IR, et al. The REFLECT statement: reporting guidelines for randomized controlled trials in livestock and food safety: explanation and elaboration. Journal of food protection. 2010;73(3):579–603.

5. Schabenberger O, et al. Introducing the GLIMMIX procedure for generalized linear mixed models. SUGI 30 Proceedings. 2005;196.

6. Schunicht OC, Guichon PT, Booker CW, Jim GK, Wildman BK, Hill BW, et al. A comparison of prophylactic efficacy of tilmicosin and a new formulation of oxytetracycline in feedlot calves. The Canadian Veterinary Journal. 2002;43(5):355.

7. White BJ, Amrine D, Goehl D. Determination of value of bovine respiratory disease control using a remote early disease identification system compared with conventional methods of metaphylaxis and visual observations. Journal of animal science. 2015;93(8):4115–4122.

8. Higgins JP, Green S. Cochrane handbook for systematic reviews of interventions. vol. 4. John Wiley & Sons; 2011.

9. Hedges LV, Olkin I. Statistical methods for meta-analysis. Academic press; 2014.

10. Borenstein M, Hedges LV, Higgins JP, Rothstein HR. Introduction to meta-analysis. John Wiley & Sons; 2011.

11. Daniels HE. Saddlepoint approximations in statistics. The Annals of Mathematical Statistics. 1954; p. 631–650.

12. Barndorff-Nielsen O, Cox DR. Edgeworth and saddle-point approximations with statistical applications. Journal of the Royal Statistical Society: Series B (Methodological). 1979;41(3):279–299.

13. Knudson C. An Introduction to Model-Fitting with the R package glmm. 2018;.

14. Wolfinger R, O’connell M. Generalized linear mixed models a pseudo-likelihood approach. Journal of statistical Computation and Simulation. 1993;48(3-4):233–243.

